# Collective MAPK Signaling Dynamics Coordinates Epithelial Homeostasis

**DOI:** 10.1101/826917

**Authors:** Timothy J. Aikin, Amy F. Peterson, Michael J. Pokrass, Helen R. Clark, Sergi Regot

## Abstract

Epithelial tissues are constantly challenged by individual cell fate decisions while maintaining barrier function. During oncogenesis, mutant and normal cells also differ in their signaling states and cellular behaviors creating competitive interactions that are poorly understood. Here we show that the temporal patterns of MAPK activity are decoded by the ADAM17-EGFR paracrine signaling axis to coordinate migration of neighboring cells and promote extrusion of aberrantly-signaling cells. Concurrently, neighboring cells increase proliferation to maintain cell density while oncogene expressing cells undergo cell cycle arrest. Moreover, the stress MAPK p38 elicits the same paracrine signaling and extrusion response, suggesting that the ADAM17-EGFR pathway constitutes a quality control mechanism to eliminate and replace unfit cells from epithelial tissues. Overall, we show that the temporal patterns of MAPK activity coordinates both single and collective cell behaviors to maintain tissue homeostasis.

## INTRODUCTION

The Receptor-Tyrosine Kinase (RTK)/RAS/ERK signaling axis (Fig. 1A) is mutated in most human cancers^1^. In normal conditions the ERK pathway promotes proliferation, differentiation, survival and cell migration^2^. Conversely, during oncogenesis mutations or amplification of ERK pathway components can also promote oncogene-induced senescence^3^ (OIS) or cellular extrusion from epithelial monolayers^4,5^. The mechanisms underlying dose dependent effects of ERK signaling have been intensely studied using bulk cell population assays. However, the advent of single cell analysis has shown that single cells often behave qualitatively different than bulk populations. In fact, *in vivo* and *in vitro* studies have now shown that pulsatile or sustained ERK activity have different effects on cell fate^6-12^. Whether oncogenic perturbations also have different functional outcomes depending on downstream signaling dynamics remains unknown. To address this question, an isogenic single-cell approach with temporal control of oncogene expression is needed.

**Fig. 1.**
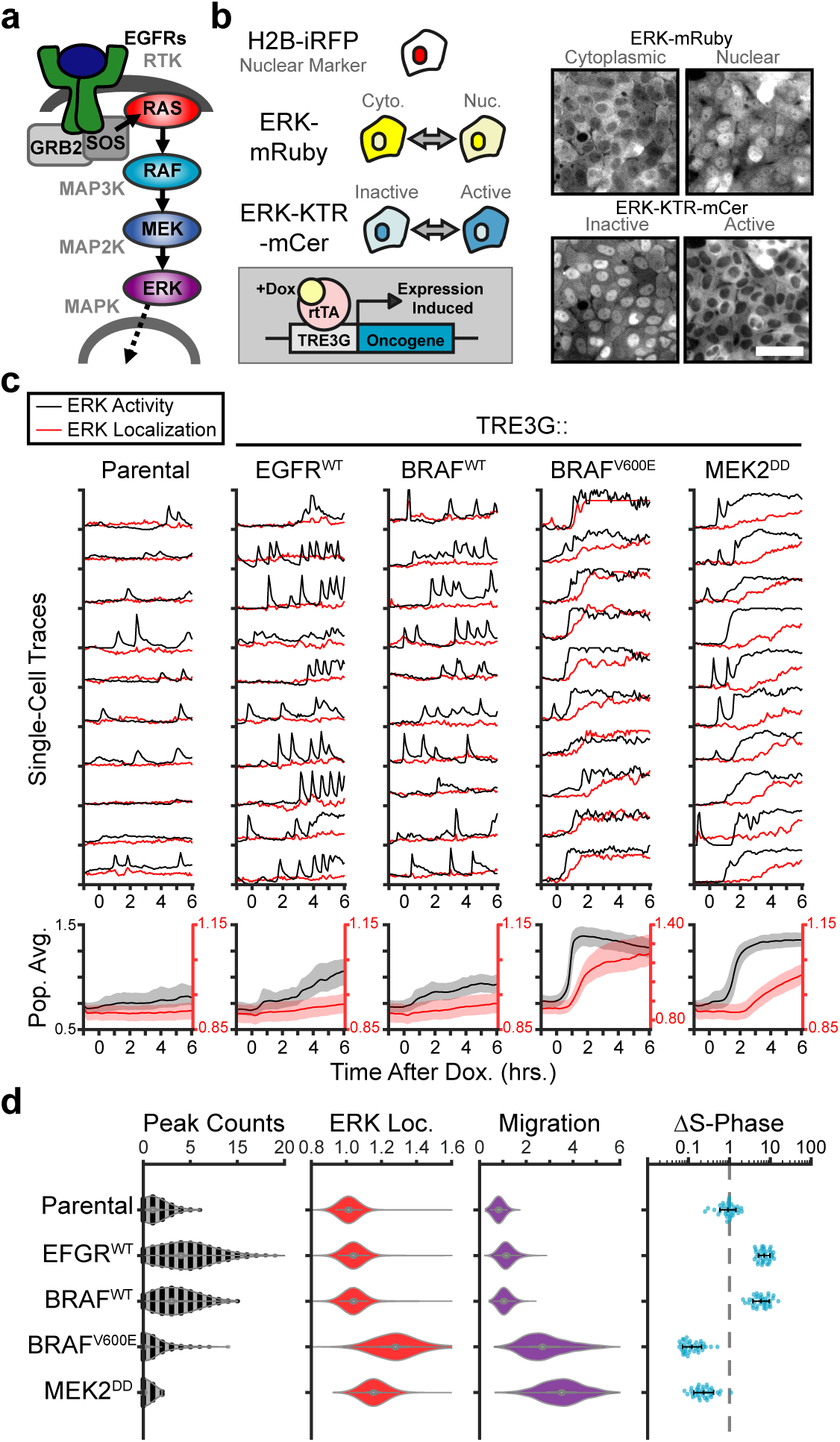
Oncogenic ERK signaling dynamics promote qualitatively different cell fates. **A**. Schematic representation of the RTK/RAS/ERK signaling pathway. **B**. MCF10A cells were transduced with lentiviral vectors expressing ERK KTR-mCerulean3 and ERK-mRuby2. The doxycycline inducible system (rTtA and TRE3G) was used to drive the expression of oncogenes during live imaging. Representative images of cytoplasmic and nuclear ERK-mRuby2 (top) and inactive or active ERK as reported by ERK KTR-mCerulean3 (bottom). Scale bar = 50µm. **C**. Cells described in B with indicated inducible oncogenes were imaged every 5 minutes for 6 hours upon doxycycline induction (Dox. 2µg/ml) at t=0. Single cells were analyzed as described in methods. Population averages represent more than 1000 cells per condition. Shaded regions indicate the 25th-75th percentiles. **D**. Quantification of data obtained in C. Single-cell counts of ERK activity peaks after induction (6-12h), ERK kinase localization fold change (final N/C ratio over basal N/C ratio per cell) and, cell migration (final over basal distance traveled per cell) were extracted as described in methods. For proliferation analysis the fraction of S-phase cells was measured using Edu incorporation and the change over the no dox control was calculated and normalized to the mean of parental cells (dashed line) (see methods). Data represents 36 independent observations.

Recent *in vivo* studies revealed that oncogene expression can trigger cell-cell competition and tissue level responses involving normal neighboring cells^13-16^. During these events, the mechanistic basis of competition and the signaling events involved in recognition between normal and diseased cells are poorly understood. Coincidentally, propagating ERK signaling waves that depend on the sheddase ADAM17 have been observed in mouse epidermis and intestinal organoids, but the physiological role of these collective signaling events in homeostasis remains unclear^7,8,17^. Understanding of the context-dependent mechanisms of paracrine signaling and cell-cell competition upon oncogene expression holds the key to unlocking new therapeutic strategies.

Here we combine live cell imaging of signaling biosensors with inducible expression of oncogenes to study the cell autonomous and non-cell autonomous effects of oncogene expression in epithelial monolayers. Our data shows that pulsatile or sustained ERK signaling resulting from oncogenic perturbations triggers different dynamics-dependent cell fates including oncogene-induced paracrine signaling via the ADAM17-EGFR signaling axis. The resulting localized signaling events are required to coordinate neighboring cell migration and active extrusion of aberrantly-signaling cells. In addition, we show that this mechanism mediates extrusion of damaged and stressed cells, highlighting the general role of MAPK signaling dynamics in coordinating individual cell fates with collective cell behavior.

## RESULTS

To study the cell autonomous and non-cell autonomous effects of oncogene expression in epithelial monolayers, we implemented a system to measure signaling dynamics in single cells during the onset of oncogene expression (Fig. 1B). Specifically, we generated monolayers derived from the chromosomally-normal human breast epithelial cell line MCF10A expressing the ERK Kinase Translocation Reporter^18^ (ERK KTR) and a fluorescently tagged ERK kinase (ERK-mRuby2). This reporter combination allowed independent measurement of ERK activity and ERK localization in live single cells at high temporal resolution. We then introduced 12 different doxycycline-inducible oncogenic perturbations via lentiviral infection and measured ERK dynamics during induction. Our results revealed two qualitatively different responses to oncogene induction: (i) increased frequency of ERK activity pulses with no change in ERK kinase localization (i.e. EGFR, B-Raf), and (ii) sustained ERK activity with subsequent nuclear translocation of ERK kinase (i.e. B-Raf^V600E^, MEK2^DD^) (Fig. 1C, Fig. S1, and Supplementary Video 1). Expression of wild-type MEK1/2 lead to gradual nuclear localization of ERK kinase without ERK activation, consistent with a role for MEK1/2 in active ERK nuclear export^19^ (Fig. S1). These data show that ERK activity and ERK localization are not always correlated, instead ERK nuclear accumulation occurs when ERK activity is maintained longer than the normal duration of ERK activity pulses.

Moreover, the two distinct dynamic patterns resulted in two qualitatively different cell fates: (i) increasing ERK activity pulses (with EGFR and B-Raf) promoted cell cycle progression^6^ while (ii) sustained ERK activity with ERK nuclear translocation (with B-Raf^V600E^ and MEK2^DD^) caused cell cycle arrest and increased migration (Fig. 1D). Importantly, these differences in cell fate where observed regardless of the point in the cascade that perturbations were introduced (i.e. both EGFR and B-Raf leading to pulsatile ERK signals) suggesting that ERK dynamics are ultimately responsible for these differences. In addition, oncogenes leading to sustained ERK activation showed different signaling onset times which correlated with the severity of migratory behaviors (Fig. S1). In particular, KRAS^G12V^ induction showed gradual activation of ERK with a moderate increase in cell migration in the first 8 hours after induction, whereas B-Raf^V600E^ led to rapid ERK activation and faster migration. Together, these data suggest that the diversity of cellular phenotypes resulting from oncogene expression depend on ERK activity dynamics.

To examine the non-cell autonomous effects of oncogene expression in epithelial monolayers, we cocultured “inducible” cells (expressing doxycycline-inducible oncogenes, a constitutively expressed H2B-mClover, and the ERK activity biosensor) with “neighboring” reporter cells (expressing MAPK live cell biosensors without inducible oncogenes) (Fig. 2A). Interestingly, expression of oncogenes that triggered sustained, but not pulsatile, ERK activity promoted ERK signaling waves in the neighboring cells (Fig. 2B-C, Fig. S2 and Supplementary Video 2). No JNK or p38 activation was observed during the first 24 hours after induction (Fig. S2). Previous studies demonstrated that ERK waves depend on the sheddase ADAM17 (Fig. 2D). which is responsible for shedding of growth factors such as HB-EGF, TGF-α, epiregulin, and amphiregulin^20^. Thus, to test the role of ADAM17 in oncogene-induced paracrine signaling, we generated a knockout (KO) cell line using CRISPR (Fig. 2E). Our results indicated that ADAM17 is necessary in inducible cells, but not neighboring cells, to trigger ERK waves in the monolayer (Fig. 2F-G and Supplementary Video 2). To identify ADAM17-released factors upon oncogene-induced paracrine signaling, we used Tandem-Mass-Tag Mass Spectrometry of supernatant proteins following induction of sustained ERK activity in WT and ADAM17^KO^ cells. A variety of known and unknown ADAM17 substrates were cleaved^20,21^, including immune surveillance proteins (HLA-A/B/C), Delta-Notch (JAG1), and Wnt (SFRP) signaling proteins (Fig. 2H and Supplementary Table 1). Of note, the EGFR ligand amphiregulin (AREG) was the most upregulated, ADAM17-dependent protein in the supernatant, supporting the idea that oncogene-induced ERK waves depend on AREG-EGFR signaling. Moreover, the inhibition of AREG receptors (EGFRs) prevented neighboring cells from activating ERK waves but did not affect ERK activity in inducible cells (Fig. 2I). Therefore, the ADAM17–EGFR signaling axis decodes ERK activity dynamics to promote ERK signaling in neighboring cells.

**Fig. 2.**
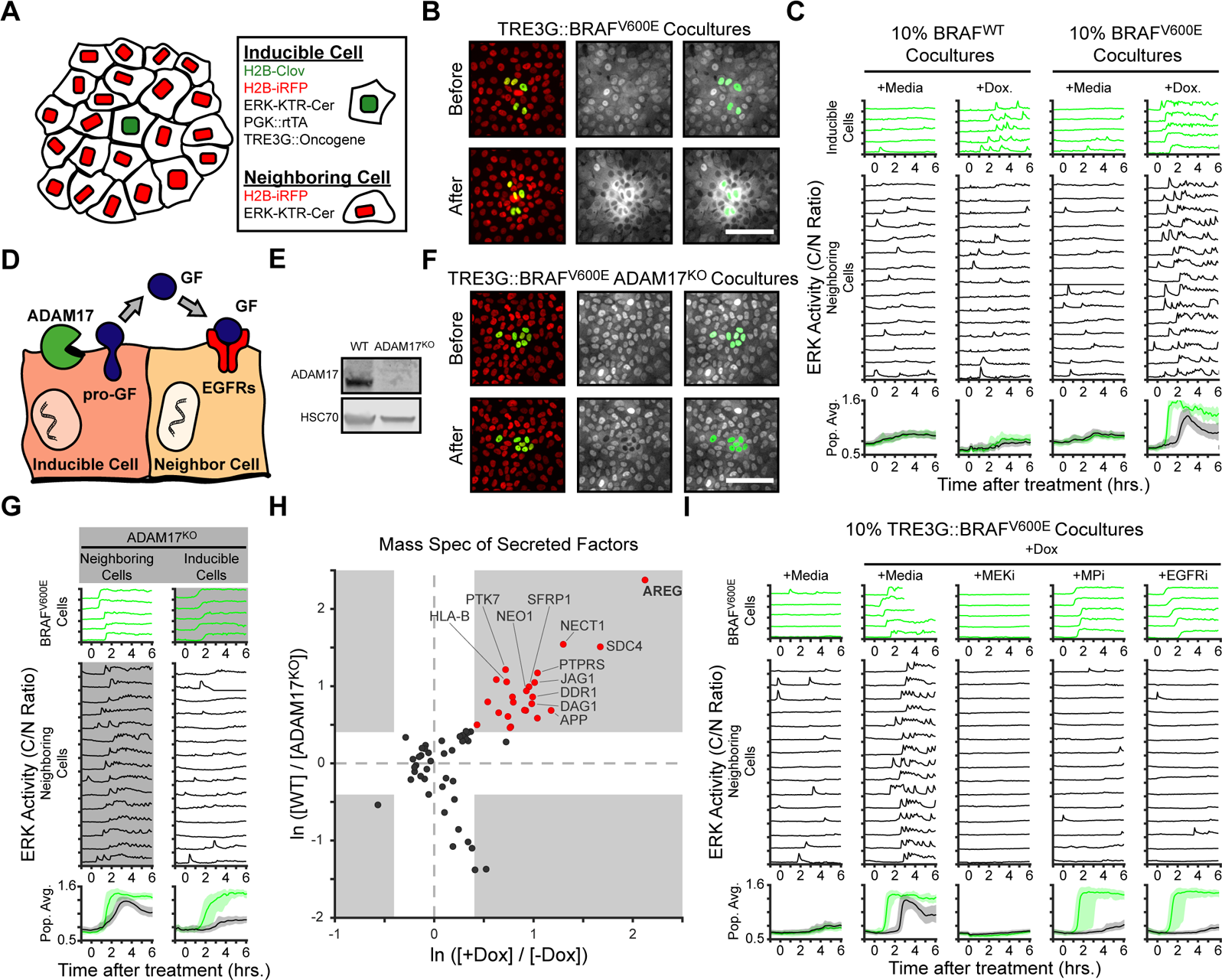
ADAM17 decodes ERK signaling dynamics to trigger ERK activity waves through the neighboring epithelium. **A**. Schematic representation of coculture assay. H2B-iRFP (red) and ERK KTR are expressed in all cells for segmentation and quantification. H2B-mClover (green) was used to label inducible cells. **B**. 10% of BRAF^V600E^ inducible cells were cocultured with ERK KTR cells and treated with doxycycline (2µg/ml). Representative images are shown. Scale bar = 100µm. **C**. BRAF^WT^ or BRAF^V600E^ inducible cells were cocultured as in B and treated with vehicle (+Media) or with doxycycline (+Dox, 2µg/ml). Single cells were quantified as described in methods. ERK activity traces in inducible (top, green) and neighboring cells (bottom, black) are shown. Population averages and 25th-75th percentiles (shaded) are shown for n > 450 cells per coculture condition. **D**. Schematic representation of the ADAM17-EGFR paracrine signaling axis. **E**. Immunoblot against ADAM17 and HSC70 in WT and ADAM17^KO^ cells (see methods for details). **F**. Representative images of ADAM17^KO^ BRAF^V600E^ inducible cells cocultured and treated as in B. **G.** ADAM17^KO^ (gray boxed traces) cells were used as inducible cells (right) or neighboring cells (left) in cocultures. Data for n > 1100 cells is presented as in C. **H**. ADAM17 substrates profiled by TMT mass spectrometry. Supernatants from ADAM17^KO^ or WT cells expressing (+Dox) or not (-Dox) BRAF^V600E^ were collected and analyzed by Tandem-Mass-Tag (TMT) mass spectrometry as described in methods. Scatter plots show the natural log of fold change values of all statistically significant (p<0.05) proteins in both WT vs. ADAM17^KO^ and +Dox vs. -Dox comparisons. Grey boxes indicate >1.5 fold change. **I**. BRAF^V600E^ co-cultured monolayers were plated as in C and pretreated with indicated inhibitors (MEKi, 5uM PD0325901; MPi, 5uM Batimastat; EGFRi, 5uM Gefitinib) for one hour. Representative single cell traces and population averages for n > 1000 cells are shown as in C.

We next addressed the physiological role of oncogene-induced paracrine signaling. Previous studies demonstrated that ERK waves orient collective cell migration during wound healing^22^. Indeed, upon oncogene expression, neighboring cell migration angles are strongly biased towards inducible cells (Fig. 3A). This effect depended on both EGFR signaling and ADAM17, suggesting that neighboring cells orient migration via a local EGFR ligand signaling gradient established by ADAM17 sheddase activity in BRAF^V600E^-expressing cells. We hypothesized that as a result of the collective migration towards shedding cells, neighboring cells could contribute to oncogenic cell extrusion^4^. In fact, confocal Z stacks of cocultures captured apical extrusion of B-Raf^V600E^ expressing cells from the monolayer (Fig. 3B and Supplementary Video 3). Expression of oncogenes that elicit cell autonomous ERK pulses without paracrine ERK waves (EGFR and B-Raf) failed to extrude after 24 hours while oncogenes that trigger sustained ERK signaling and ERK waves in the neighboring cells (B-Raf^V600E^ and MEK2^DD^) were efficiently extruded (Fig. 3C).

**Fig. 3.**
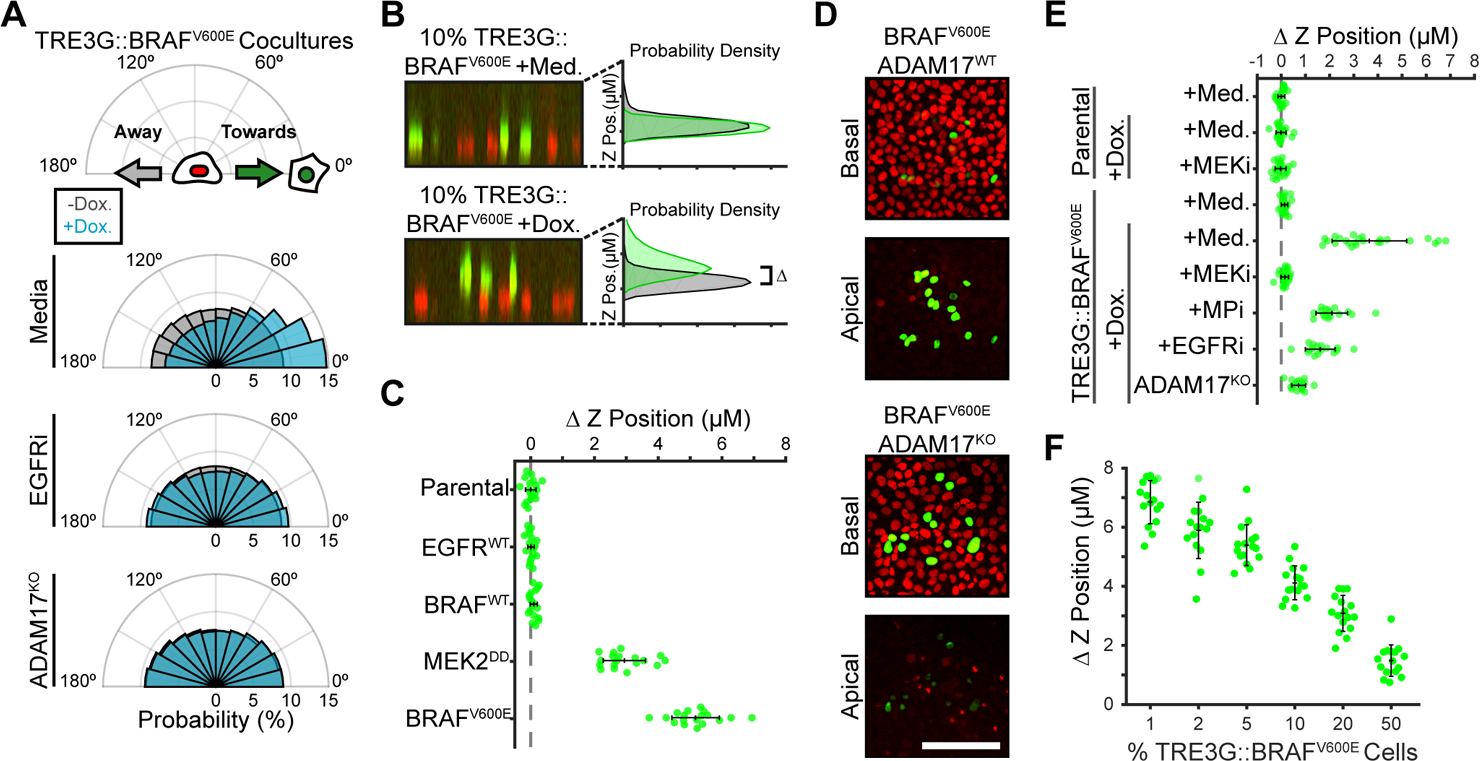
Paracrine ERK activation coordinates extrusion of aberrantly signaling cells by the neighboring epithelium. **A**. BRAF^V600E^ inducing cells (WT or ADAM17^KO^) were plated in 1% cocultures and treated with doxycycline (2µg/ml) in the presence or absence of EGFR inhibitor gefitinib (5µM) as indicated. Radial histograms represent angle distributions before (grey) and after (cyan) induction. Data represents n > 1000 cells from 10 independent observations per condition. **B.** 10% BRAF^V600E^ Cocultured monolayers were seeded and starved as described in methods. After 24 hours with doxycycline (2µg/ml), monolayers were imaged by spinning disk confocal. Representative orthogonal Z projections and probability densities of inducible (green) and neighboring (grey) cells are shown (see methods). Extrusion (ΔZ) is calculated as the difference between the gaussian fitted maxima between green and black distributions. **C**. 10% cocultures of indicated parental or inducible cells were plated and imaged as in B. Data represents 17 independent observations normalized to the mean of parental cells (dashed line) with mean and +/- standard deviation (black bars). **D.** Representative basal and apical images (+6 µM) of WT or ADAM17^KO^, BRAF^V600E^ inducible cells (green) in WT monolayers (red) after 24h of doxycycline treatment. **E**. Distributions of relative z-positions for indicated cocultures pretreated with inhibitors (MEKi, 5µM PD0325901; MPi, 5µM Batimastat; EGFRi, 5µM Gefitinib), imaged and analyzed as in B for n > 16 observations. **F**. Inducible BRAF^V600E^ cocultures at indicated proportions were plated, treated and analyzed as in B. n > 15 observations per condition.

KRAS^G12V^ induction did not result in extrusion as previously observed in MDCK cells^4^. However, since sustained ERK activation in KRAS^G12V^ occurs later than B-Raf^V600E^ (Fig. S1) extrusion may also occur at a later time. Critically, extrusion of B-Raf^V600E^ expressing cells was abolished in ADAM17^KO^ cells or in the presence of EGFR inhibitor (Fig. 3D-E). This result suggested that activation of inducible cells alone is not sufficient for extrusion and that neighboring cell EGFR-ERK activation is required. In addition, extrusion of B-Raf^V600E^ expressing cells was less efficient with increasing proportions of inducible cells, reinforcing the idea that the coordinated behaviors of neighboring cells depend on signaling gradients that are better established with fewer sources of signaling factors (Fig. 3F). We also found that other genes known to trigger epithelial cell extrusion^23^ such as Cdc42^G12V^ similarly trigger ERK signaling waves in neighboring cells (Fig. S3). Taken together, our data indicates that EGFR and ADAM17-dependent ERK signaling waves in neighboring cells are necessary for oncogenic cell extrusion.

The frequency of ERK activity pulses promotes cell proliferation^6^ and neighboring cells surrounding B-Raf^V600E^ expressing cells showed highly pulsatile ERK signaling (Fig. 2C,I). Thus, we asked whether oncogene expression promotes cell proliferation in a non-cell autonomous way. Strikingly, while sustained ERK signaling caused cell autonomous cell cycle arrest (Fig. 1D), neighboring cell proliferation increased by 5-8 fold (Fig. 4A-B). These data indicate that, depending on ERK activity dynamics, oncogene expressing cells can promote either cell autonomous or non-cell autonomous tissue growth. Importantly, the non-cell autonomous contributions depend on ADAM17 activity which can be targeted with existing inhibitors and monoclonal antibodies^20,21^.

**Fig. 4.**
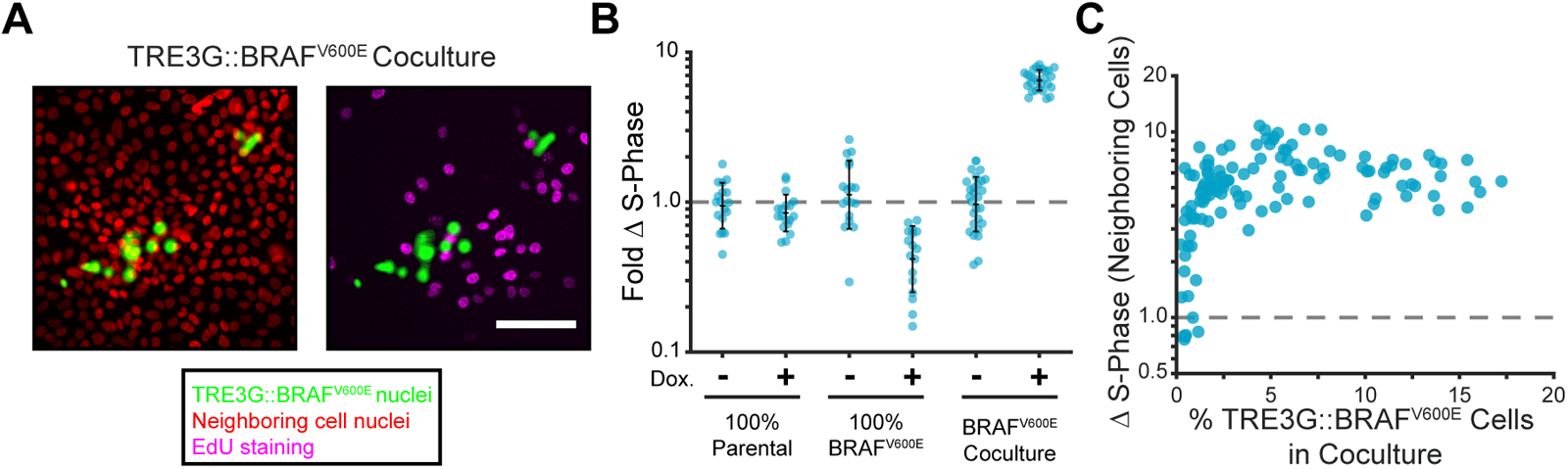
Oncogene-induced paracrine signaling promotes proliferation of surrounding cells. **A**. Representative images of BRAF^V600E^ cocultures treated with doxycycline and EdU as described in methods. Inducible cell marker (green), all nuclei (red) and edu staining (magenta) are shown. Scale bar = 100µm. **B**. Indicated monolayers were cultured and incubated with or without doxycycline for 24 hours. The change in S phase cell fractions was determined by EdU incorporation as described in methods and normalized to parental mean (dashed line). Bar represents mean and standard deviation for n > 26 observations. **C**. Inducible BRAF^V600E^ cocultures were plated at different proportions and labelled with EdU as in A. The fold-change in S-phase cell fractions is plotted against the percent of BRAF^V600E^-expressing cells for each position. 123 total observations shown.

Apoptotic cells are commonly extruded from epithelial monolayers^24^. In fact, we observed ERK waves associated with spontaneous cell death events in our live imaging experiments (Supplementary Video 4). To determine whether the ADAM17-EGFR signaling axis plays a role in extrusion during cell death, we used a digital mirror device to stimulate specific regions of monolayers with UV light. Using ERK and p38 live-cell biosensors, we observed synchronous pulses of ERK and p38 activity in irradiated cells followed by ERK signaling waves that traveled 3-4 cell diameters through the surrounding monolayer (Fig. 5A-C, Fig. S4 and Supplementary Video 5). UV-induced ERK signaling waves were dependent on ADAM17 and EGFR signaling and demonstrated a rigorous temporal correlation with cell migration towards the irradiated cells (Fig. 5D-F).

**Fig. 5.**
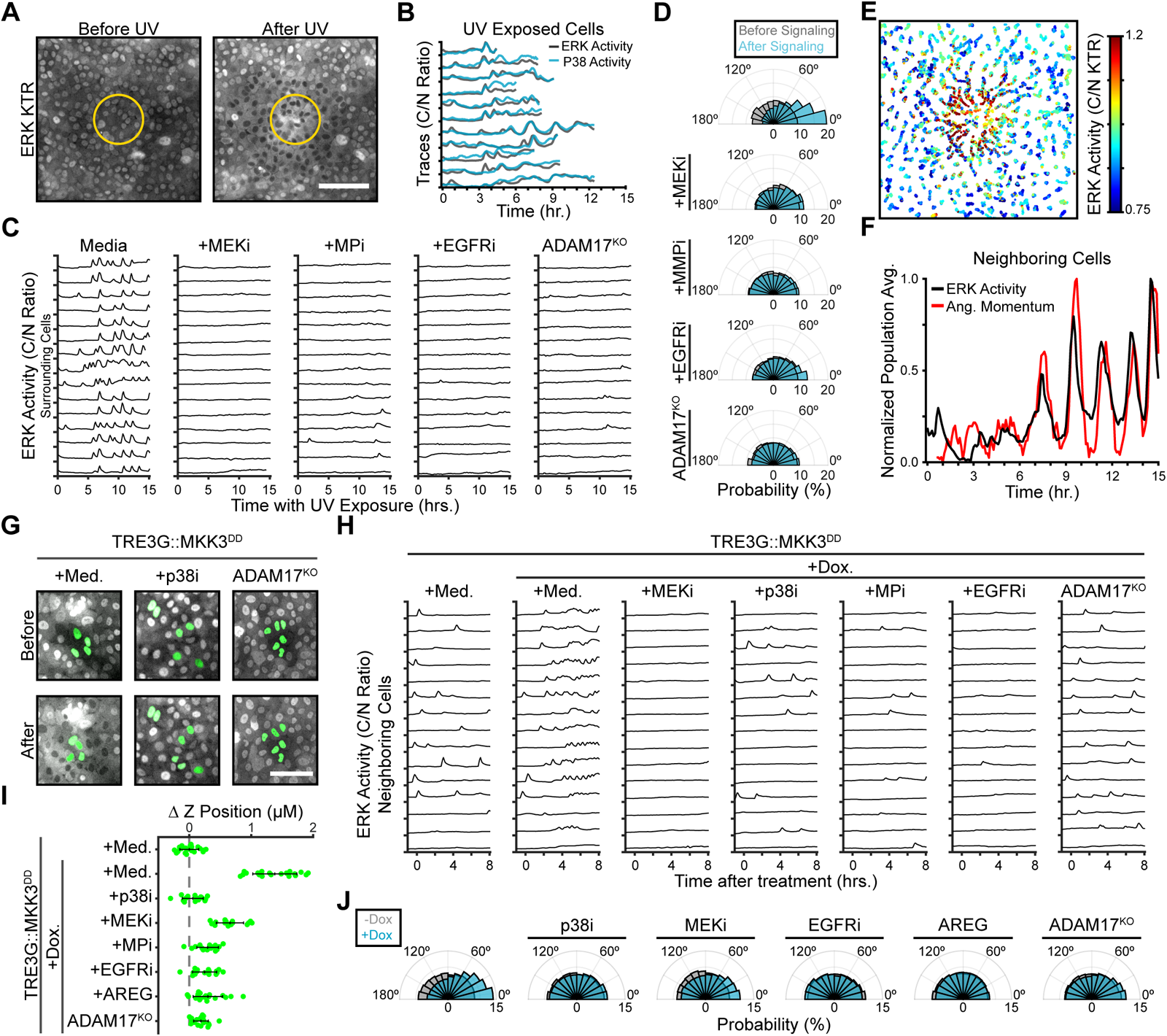
ADAM17-EGFR dependent ERK activation coordinates extrusion in response to UV exposure and p38 activation. **A**. Representative ERK KTR images of a MCF10A monolayer before and after UVA irradiation (365nm) within the yellow circle. Scale bar = 100µm. **B**. MCF10A cells expressing ERK and p38 KTRs were imaged during pulsed UVA light (365nm) exposure of a defined area using a digital mirror device (DMD). C/N ratios of ERK-KTR (grey) and p38 KTR (cyan) sensors within the illuminated region are shown. **C**. WT or ADAM17^KO^ cells were UV irradiated as in B and pretreated with indicated inhibitors (MEKi, 5uM PD-0325901; MPi, 5uM Batimastat; EGFRi, 5uM Gefitinib). Representative ERK activity traces of cells outside the exposed area are shown. **D**. Migration angles were calculated for cells imaged and quantified in C. Radial histograms represent directed migration before (grey) and after (6-12h, cyan) ERK waves. Data represents n > 180 cells from 2 observations per condition. **E**. Data obtained in panel D is represented as a scatter plot of nuclear positions of each cell over time. Colormap represents C/N ERK KTR ratio. **F**. ERK activity (black) and mean angular momentum (red) data for all cells outside of the UV-exposed region towards the exposed region in F. See methods for details. **G**. WT or ADAM17^KO^ cells with inducible MKK3^DD^ (green) were cocultured with neighboring cells expressing ERK-KTR (grey). p38 inhibitor (5µM BIRB-796) was added when indicated. Scale bar = 100µm. **H**. MKK3^DD^ inducible cells were plated in 10% coculture, treated and imaged as in 2B. 15 representative neighboring cell traces are shown for each condition. **I**. Radial histograms of migration angles for cells obtained in D before (grey) and after (cyan) induction presented as in 3A. Data represents n > 900 cells from > 6 observations per condition. **J**. Inducible cocultured cells were plated as in H and treated with indicated inhibitors. Extrusion was calculated as in 3B-C and described in methods. Bars represent mean and standard deviation for n > 16 observations.

The stress MAPK p38 has been previously shown to activate ADAM17 shedding^25^. Thus, to separate the contributions of ERK and p38 in regulating ADAM17 activity, migration, and extrusion, we used doxycycline inducible expression of MKK3^DD^, a constitutively-active MAP2K specific for p38 (Fig. S5). With this system, p38 activation led to ERK signaling waves in neighboring cells that depended on ADAM17 and EGFR signaling (Fig. 5G-H and Supplementary Video 6). Of note, B-RAF^V600E^-induced paracrine signaling was not abolished by p38 inhibition, indicating that sustained ERK activation and p38 activation are each capable of activating ADAM17 (Fig. S6). MKK3^DD^ cocultures also allowed us to induce ADAM17 shedding in an ERK-independent manner and directly test the role of ERK activity in neighboring cells to promote extrusion of shedding cells. Inhibiting either ERK or EGFR activity abolished directed migration and extrusion of p38-active shedding cells (Fig. 5I-J) and Fig. S7). Additionally, exogenous AREG disrupted the coordination of ERK signaling waves, preventing directed migration of neighboring cells and extrusion of shedding cells (Fig. 5I-J). These data suggest that local paracrine signals are essential to coordinate extrusion, limiting the value of conditioned media experiments to study this process. In fact, cell competition studies have proposed that short range diffusible signals differentiate winner and looser cell populations^15,16,26^. Other studies have shown that sphingosine-1-phosphate (S1P) production plays an important role in epithelial extrusion^27-29^. While inhibition of sphingosine kinases (SphKs) did not abolish ERK signaling waves, the effect on extrusion was not additive with ADAM17 ablation or EGFR inhibition (Fig. S8), indicating that the role of S1P in extrusion may be linked to its crosstalk with EGFR-ERK signaling. Indeed, ERK can stimulate production of S1P by activating Sphingosine Kinases^30,31^ and S1P stimulates EGFR activation during development and in glioblastomas^32,33^. Taken together these data demonstrate that either ERK or p38 MAPK signaling dynamics can initiate ADAM17-EGFR paracrine signaling to promote extrusion of unfit cells from monolayers, therefore coordinating epithelial homeostasis.

## DISCUSSION

The mutation occurrence in different tumor types is robustly biased by often unclear mechanisms ^1,34,35^. To understand context dependent effects of specific oncogenic mutations, quantitative, live single cell approaches offer precise spatial and temporal resolution which enables dissection of cell autonomous and non-cell autonomous effects in real time. In addition, given the species - specific differences in tumorigenesis, the use of cultured, non-transformed human cells can be more revealing for human disease than mouse models. Using a live single cell approach in human mammary epithelial cells, we have shown that different ERK pathway oncogenes cause unique temporal patterns of ERK activity. These dynamics control a variety of phenotypes including migration, proliferation, cell cycle arrest, and oncogene-dependent paracrine signaling. The dramatic difference in cell fate based solely on dynamics suggests that cell-type dependent extrinsic factors which modify signal transduction may contribute to the biased oncogene occurrence of different tumor types. In addition, this data highlights the importance of quantitative live single cell approaches to understand the effects of genetic perturbations.

Moreover, we show that oncogene-induced paracrine signaling occurs before the onset of extrusion and only when ERK activity is sustained (upon expression of either BRaf^V600E^ or MEK2^DD^) indicating that the ADAM17-EGFR signaling axis can decode temporal patterns of ERK activity. Thus, aberrantly-signaling cells are recognized by neighboring cells via ectodomain cleavage of signaling factors. In the case of mammary epithelia, we show that AREG shedding by aberrantly-signaling cells generates localized EGFR signaling (Fig. 2H-I) that coordinates migration of neighboring cells to promote extrusion (Fig. 3). This finding agrees with previous work highlighting the role of AREG in mammary epithelial homeostasis^36^. In addition, our mass spec results also show that shedding interferes with antigen presentation, suggesting that ADAM17 activity could interfere with cancer immunotherapies ^21,37^. The connection between shedding, extrusion and immune surveillance proposed here provides a new framework to understand cell-cell competition upon expression of oncogenes. Future studies will determine the roles of shedding and extrusion in cancer progression *in vivo*.

Cell competition occurs in tissues when differential fitness results in elimination of “loser” cells and expansion of “winner” cells^26,38^. However, little is known about the way in which unfit cells are recognized by their neighbors before elimination. In drosophila, competitive recognition can involve secretion of JNK and NF-κB stimuli or juxtracrine comparison of cell surface proteins^39^. In mammalian tissues, high Myc cells force apoptosis, entosis, or differentiation of neighboring cells depending on the context^14-16^. We propose that the regulation of the ADAM17-EGFR paracrine signaling axis by MAPK signaling dynamics contributes to the recognition of distinct cell populations within tissues that leads to cellular competition.

In addition to oncogene expression, UV exposure and activation of the stress MAPK p38 leads to an identical paracrine signaling and extrusion response. This suggests that the ADAM17-EGFR pathway represents a general mechanism to replace unfit cells from epithelial monolayers. Using p38-dependent shedding we demonstrated that neighboring cell ERK activation is required to extrude aberrantly-signaling cells. However, in response to oncogenes, the difference between extruded and neighboring cells is the dynamics of ERK signaling (pulsatile vs. sustained). Thus, targeting ERK signaling with general kinase inhibitors will affect both ligand shedding and extrusion processes. We anticipate that new drugs specifically designed to alter ERK signaling dynamics will leverage cell-cell competition within tissues to improve therapeutic outcomes.

In summary, our data indicate that MAPK signaling dynamics simultaneously regulate individual cell fates and the paracrine signaling events necessary to maintain epithelial homeostasis. During oncogenesis, disrupted signaling dynamics leads to paracrine signaling and cell-cell competition that bias cell populations. We propose a critical role for signaling dynamics and the ADAM17-EGFR signaling axis in the recognition of oncogenic and stressed cells that precedes competitive elimination.

## METHODS

### Cell Lines & Reagents

MCF10A human mammary epithelial cells (ATCC) were grown at 37° and 5% CO_2_ in DMEM/F12 (Gibco) with 5% horse serum (HS) (Sigma), 10 µg/ml Insulin (Sigma), 20 ng/ml EGF (Peprotech), 1x Penicillin-Streptomycin (P/S) (Gibco), 0.5 mg/ml Hydrocortisone (Sigma), 100ng/ml Cholera Toxin (Sigma). Cells were passaged every 3 days with 0.25% Trypsin-EDTA (Gibco).

Cell lines were generated with lentivirus produced in HEK293-FTs (Thermo) with third-generation packaging plasmids. Viral supernatants were collected 48 hours after transfection and incubated in MCF10As with polybrene (10 µg/ml, EMD Millipore). To create dual-sensor cells, MCF10As were infected with a lentiviral H2B-iRFP vector (Addgene) and sorted. We used gateway cloning^40^ to introduce ERK-KTR-mCer3 and ERK1-mRuby2 into PGK pLenti DEST vectors (Addgene), infected and selected the H2B-iRFP MCF10As (Blasticidin 3µg/ml and Hygromycin 10 µg/ml Corning). We isolated moderately expressing clones using cloning cylinders (EMD Milipore). For inducible cells, a gateway-ready reverse TET trans-activator (rtTA) plasmid was created by adding the rtTA with a 2A peptide to the Puromycin resistance gene in a CMV Puro DEST plasmid (Addgene) by gibson cloning^41^. Human coding sequences were acquired from either Addgene or the Thermo Ultimate ORF Collection, sequence verified, and introduced in the rtTA CMV Puro DEST plasmid by gateway cloning^40^. These plasmids were used for lentivirus, and infected cells were selected with Puromycin (1 µg/ml, Sigma).

For inhibitor experiments, small molecules and doxycycline were dissolved to a 10X working concentration in imaging media before addition. Final DMSO concentration did not exceed 0.15%. Inhibitors used include the MEK inhibitor PD-0325901, the MMP/ADAM inhibitor Batimastat, the EGFR inhibitor Gefitinib, the p38 inhibitors BIRB-796, and the Sphingosine Kinase inhibitor SKII all from Selleck Chemicals. The p38 inhibitor SB-203580 was obtained from Sigma. Amphiregulin was ordered from Peprotech.

The ADAM17-KO cell lines were created using the CRISPR V2 Neo system (a gift from Dr. Andrew Holland) and gRNA oligos targeting R241 of exon 6. Dual sensor cells were infected with lentivirus carrying this plasmid, selected with Neomycin (500µg/ml, Sigma) and clonally expanded before western blot validation (Fig. 2B).

### Live Imaging

Cells were plated at 3*10^5^ cells/well in fibronectin-treated (EMD Millipore) 96-well glass-bottom plates (Thermo Scientific) 48 hours before imaging. The following day, monolayers were serum-starved with 0.5% HS, phenol-red-free DMEM/F12 containing P/S with 1mM Na Pyruvate and 10mM HEPES. For signaling experiments, media was switched to 0% HS several hours before imaging to limit basal signaling. Monolayers were imaged using a Metamorph-controlled Nikon Eclipse Ti-E epifluorescence microscope with a 20x air objective and a Hamamatsu sCMOS camera. The multi LED light source SpectraX (Lumencor) and the multiband dichroic mirrors DAPI/FITC/Cy3/Cy5 and CFP/YPF/mCherry (Chroma) where used for illumination and imaging without any spectral overlap. For regional UV illumination, a Digital Mirror Device (DMD, Mosaic3 from Andor) limited exposure of samples to 365nm light within a central 100µm radius every timepoint. For extrusion experiments, a Metamorph-controlled Nikon Eclipse Ti-E spinning-disc confocal (Yokogawa W1) with a 20x objective, Prime 95-B sCMOS camera (Photometrics) and a Multiline laser launch (Cairn Research) was used to capture H2B-iRFP and H2B-mClover images every 1µm of a 25-30µm range through monolayers. Temperature, humidity and CO_2_ were maintained throughout all imaging using OKO Labs control units. Sample sizes were selected by attempting to capture at least 100 cells from each population, with several hundred cells preferred. Key conditions from imaging experiments were performed at least twice, with one replicate presented in figures.

### Image Analysis and Quantification

Primary time-lapse images were subjected to flat-fielding and registration (custom software) before object segmentation and measurements in Cell Profiler. Nuclear positions were used to track individual cells through time-series (custom software) and intensity ratios were calculated as previously described^18^. Minimal cleaning of traces excluded cells where tracks switched between two objects, where the KTR ratios were affected by segmentation errors, or where traces represent less than two thirds of the entire time-course. In conditions where cells move rapidly, such as BRAF^V600E^ and MEK2^DD^, and traces are shorter due tracking errors, track-length restraints were relaxed to include more cells for analysis. Single cell traces were chosen by random plotting of distinct cells and selection of those that were tracked throughout the whole experiment. Directed migration was quantified by positional changes over 20-minute intervals for specified time windows. The migration angles of neighboring cells are plotted as radial histograms where 0° indicates migration directly towards, and 180° directly away from the center of isolated inducible cell groups (Fig. 3A and 5J) or the center of UV illumination (Fig. 5D). Angular momentum in Fig. 5G is quantified from the average direction and magnitude of cell migration towards the center of the exposed region. For extrusion experiments, histograms of mClover and mRuby pixel intensities across each z-stack were fit to gaussian curves using Matlab. The difference in gaussian fitted maxima of inducible cells and neighboring cells for each observation are plotted. Extrusion experiment sample size represents all non-overlapping positions from 2-3 independent wells excluding outliers resulting from imaging artifacts.

### Proteomics

For validation of ADAM17 CRISPR-KOs, suspected clones were lysed with RIPA buffer (CST) containing HALT protease & phosphatase inhibitors (Thermo), and reduced in Laemelli SDS buffer (BioRad) with BME (Sigma). Samples underwent electrophoresis on 4-15% gradient polyacrylamide gels (BioRad) and were immunoblotted with Rabbit anti-ADAM17 (CST) and mouse anti-HSC70 (Santa Cruz Biotechnology) primary, and IRDye donkey anti-rabbit 800 and goat anti-mouse 680 secondary antibodies (LiCor) before imaging with an Odyssey Infrared Scanner (LiCor).

For mass spectrometry, cells were grown to 90% confluency in T175 flasks and serum starved 24 hours (see live imaging) before switching to 15mL growth factor/serum-free DMEM/F12 +/- Dox for 4 hours. The supernatant was collected and concentrated using 3kDa cut-off centrifugal filters (Millipore-Sigma). Triplicate samples were quantified by the Pierce Assay (Thermo Scientific), reduced, alkylated, and trypsin digested before labeling with Tandem Mass Tag labels. Peptide fractions were analyzed by LC/MSMS using an Easy-LC 1200 HPLC system interfaced with an Orbitrap Fusion Lumos Tribrid Mass Spectrometer (Thermo Fisher Scientific). Isotopically resolved masses in precursor and fragmentation spectra were processed in Proteome Discoverer software (v2.3, Thermo Scientific). Data were searched using Mascot (2.6.2, Matrix Science) against the 2017_Refseq 83 Human database and filtered at a 1% FDR confidence threshold.

### Cell proliferation assay

Monolayers were plated and starved as described above and treated with Dox in the presence of indicated inhibitors for 24 hours. During the final 4 hours, EdU (10µM, Thermo Fischer Scientific) was added into cultures to label S phase cells then fixed with methanol and washed before Alexa-Fluor Azide 488 click labelling (Thermo Fischer Scientific) and DAPI staining (Thermo Scientific). Monolayers were imaged by epifluorescence. Because methanol fixation eliminates fluorescence from fluorescent proteins, cocultures were imaged just before fixation and registered with DAPI and EdU images to determine positions of inducible and neighboring cells. Sample size for population EdU experiments represents all non-overlapping positions from 2-3 independent wells, excluding outliers resulting from imaging artifacts. Key conditions were replicated at least twice.

## Acknowledgments

We thank all members of the Regot and Holland labs for helpful discussions and technical advice and the Mass Spectrometry and Proteomics Core at JHSOM for assistance processing samples. We acknowledge our funding sources: NIH T32 pre-doctoral training grants to T.A, A.P., M.P., and H.C. (GM007445). NSF Graduate Research Fellowships to T.A., M.P. and H.C. (DGE-1746891). An NSF CAREER award (MCB-1844994), NIGMS R35 (1R35GM133499), American Cancer Society Research Scholar Grant (133537-RSG-19-005-01-CCG), and Jerome L. Greene Foundation Discovery Award to S.R.

**The authors declare no competing interests**.

## Supplementary Video Legends

**Supplementary Video 1. | Different ERK Dynamics Following Oncogene Induction**.

Dual Sensor MCF10A cell lines containing H2B-iRFP nuclear marker (red), ERK-mRuby2 localization reporter (yellow), and an ERK KTR-mCerulean3 representing kinase activity (cyan), and indicated inducible oncogene constructs were treated with doxycycline (2µg/ml) at time 0 (see Fig. 1). Images were collected every five minutes for an hour basal period and 12 hours after induction. Scale bar = 100µm.

**Supplementary Video 2. | Oncogene-Induced ERK Signaling Waves**

Inducible BRAF^V600E^ cells with a H2B-mClover nuclear marker (green) were cocultured in monolayers with neighboring cells (H2B-iRFP, red) (see Fig. 2). Images were collected every 5 minutes for an hour basal period and 12 hours following addition of doxycycline (2µg/ml) at time 0. ERK KTR-mCer3 (grey) reports ERK activity in all cells. Three examples are shown for both WT (left) and ADAM17^KO^ (right) inducible cells. Scale bar = 100µm.

**Supplementary Video 3. | Extrusion of BRAF**^**V600E**^**-Expressing Cells**

Cocultured monolayers of inducible BRAF^V600E^ cells were imaged every 10 minutes for 24 hours following doxycycline (2µg/ml) treatment. The first frame shows the starting location of inducible cells (H2B-mClover, green) and neighboring cells (H2B-iRFP, red). The following video indicates cells in a basal position within the monolayer in red, and cells in an apical z-plane (+8µm) in cyan. Scale bar = 100µm.

**Supplementary Video 4. | ERK Signaling Waves Following Spontaneous Cell Death**

Nine examples are shown for spontaneous cell death events in serum-starved MCF10A monolayers imaged every 5 minutes. ERK KTR-mCer3 represents ERK activity (grey). Scale bar = 100µm.

**Supplementary Video 5. | UV-Induced ERK Signaling Waves**

Monolayer cultures were exposed to strong 365nm UVA light every 6 minutes for 15 hours within a central 100µm radius using a digital-mirror-device (see Fig. 4). ERK KTR-mCer3 reports ERK activity (grey) for the position shown in Fig 4a. Scale bar = 100µm.

**Supplementary Video 6. | p38-induced ERK Signaling Waves**

Inducible MKK3^DD^ cells with a H2B-mClover nuclear marker (green) were cocultured in monolayers with neighboring cells (H2B-iRFP, red) (see Fig. 4). Images were collected every 5 minutes for an hour basal period and 24 hours following addition of doxycycline (2µg/ml) at time ERK KTR-mCer3 (grey) reports ERK activity in all cells. Three examples are shown for both WT (left) and ADAM17^KO^ (right) inducible cells. Scale bar = 100µm.

**Supplementary Figure 1.**
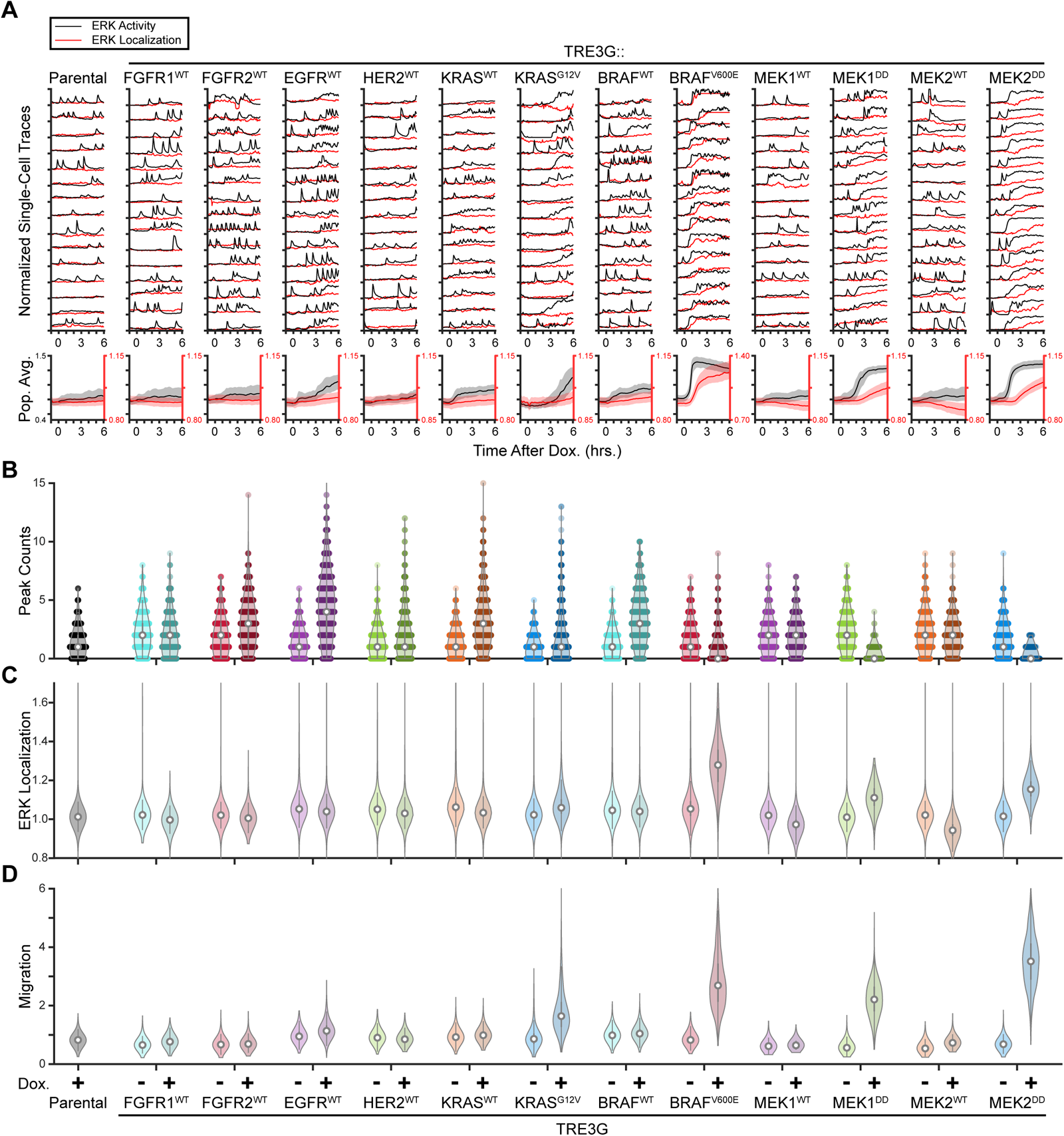
A screen for oncogenic effects on ERK dynamics and cell behavior. **A**. Inducible cells expressing indicated genes treated and plotted as in 1C. Single traces and population data are reproduced for oncogenes appearing in Fig. 1. **B-D**. Single cell peaks, ERK nuclear localization, and migration are quantified directly from imaging data as in 1D for indicated inducible cells in the presence or absence of doxycycline (2µg/ml). Data appearing in Fig. 1 is reproduced.

**Supplementary Figure 2.**
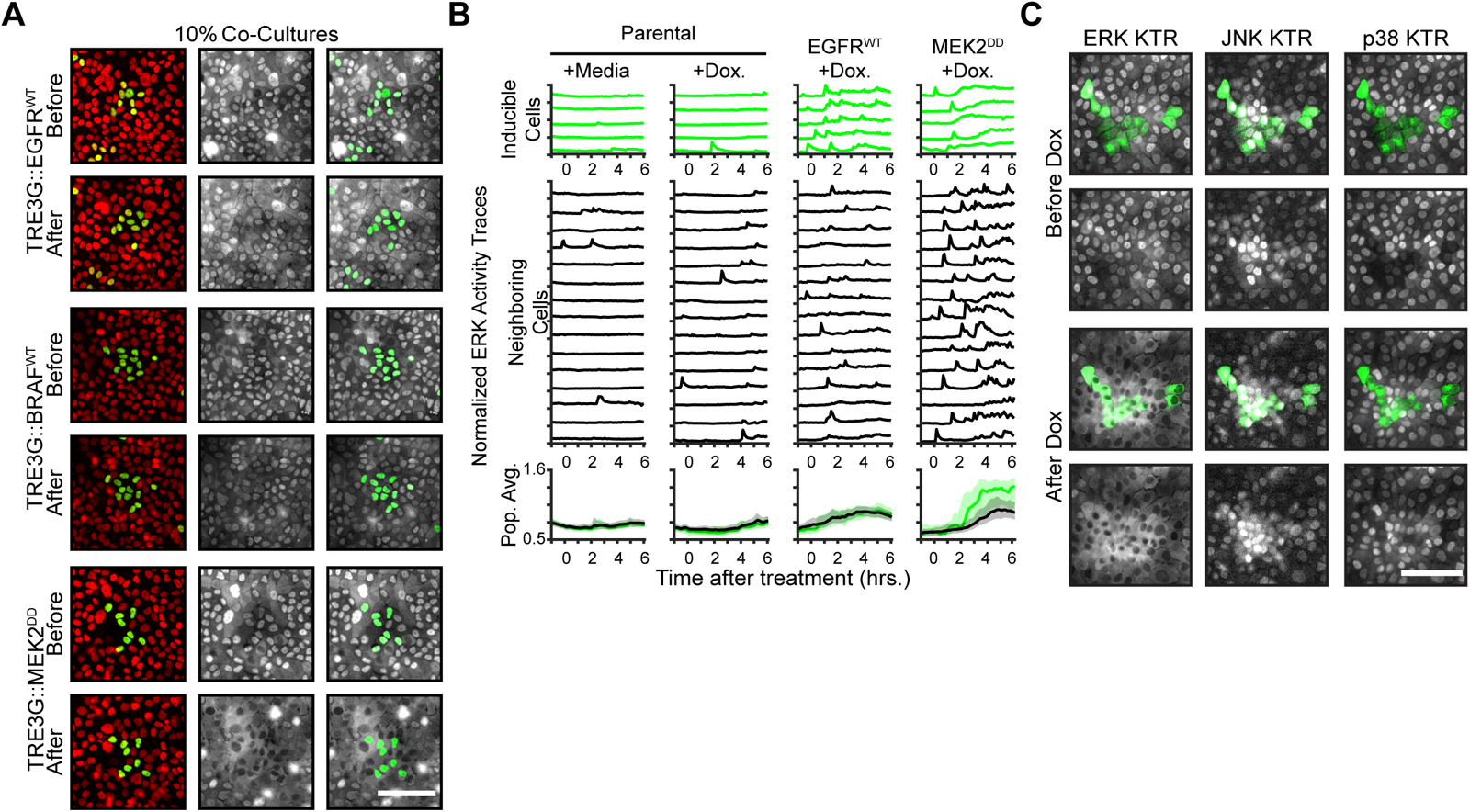
ERK dynamics-dependency of paracrine ERK activation. **A**. Images from cocultures of inducible BRAF^WT^, EGFR^WT^, and MEK2^DD^ cells presented as in 2B. Scale bar = 100µm. **B**. ERK activity traces from cocultures of indicated cells treated as in 2C. Parental cells have an H2B-mClover nuclear marker without any inducible gene system. Population averages and 25th-75th percentiles (shaded region) shown for n > 1000 cells. **C**. The TRE3G-BRAF^V600E^ construct was expressed in a cell line containing ERK KTR-mCer3, p38 KTR-mClov, and JNK KTR-mRuby2. These cells were incubated with CellTracker Deep Red dye (green, Invitrogen) and cocultured with unlabeled neighboring cells. Images display the activity of ERK, p38, and JNK before and 6 hours after addition of doxycycline (2µg/ml). Scale bar = 100µm.

**Supplementary Figure 3.**
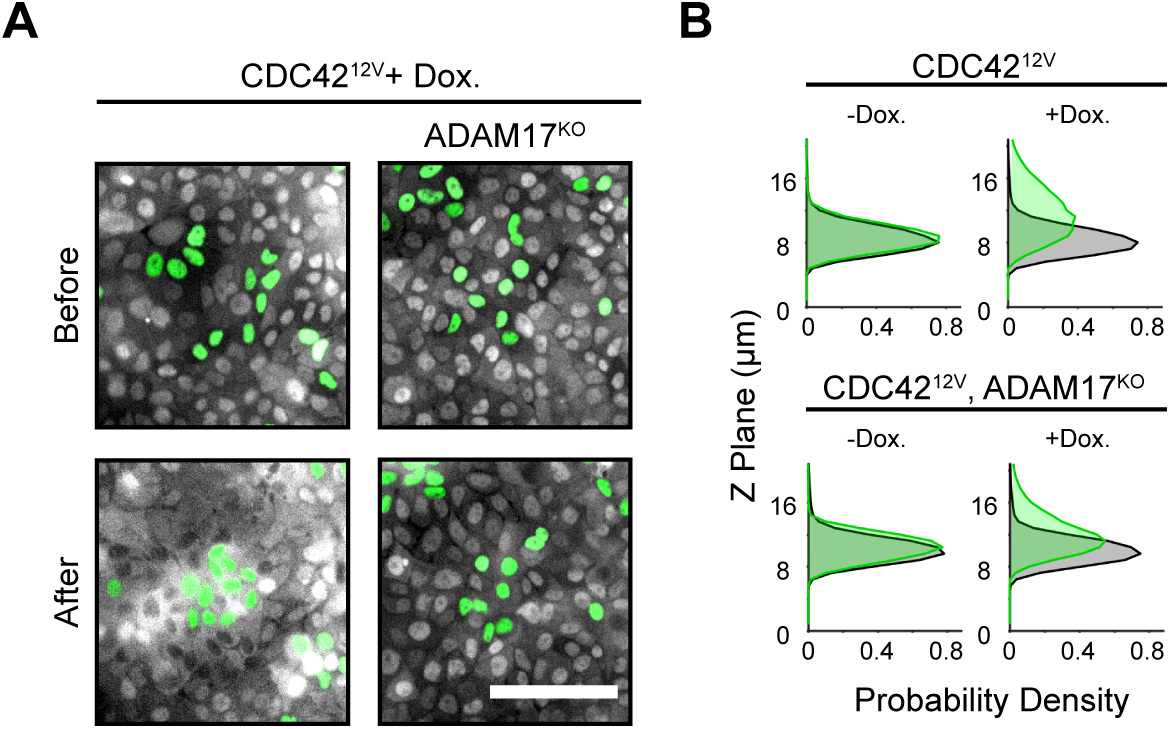
CDC42^12V^ results in paracrine ERK signaling before extrusion. **A**. Representative images from 10% cocultures of WT or ADAM17^KO^, CDC42^12V^ inducible cells plated and treated as in 2B. Scale bar = 100µm. **B**. Histograms from confocal Z stacks demonstrate extrusion of CDC42^12V^ inducible cells as in 3B.

**Supplementary Figure 4.**
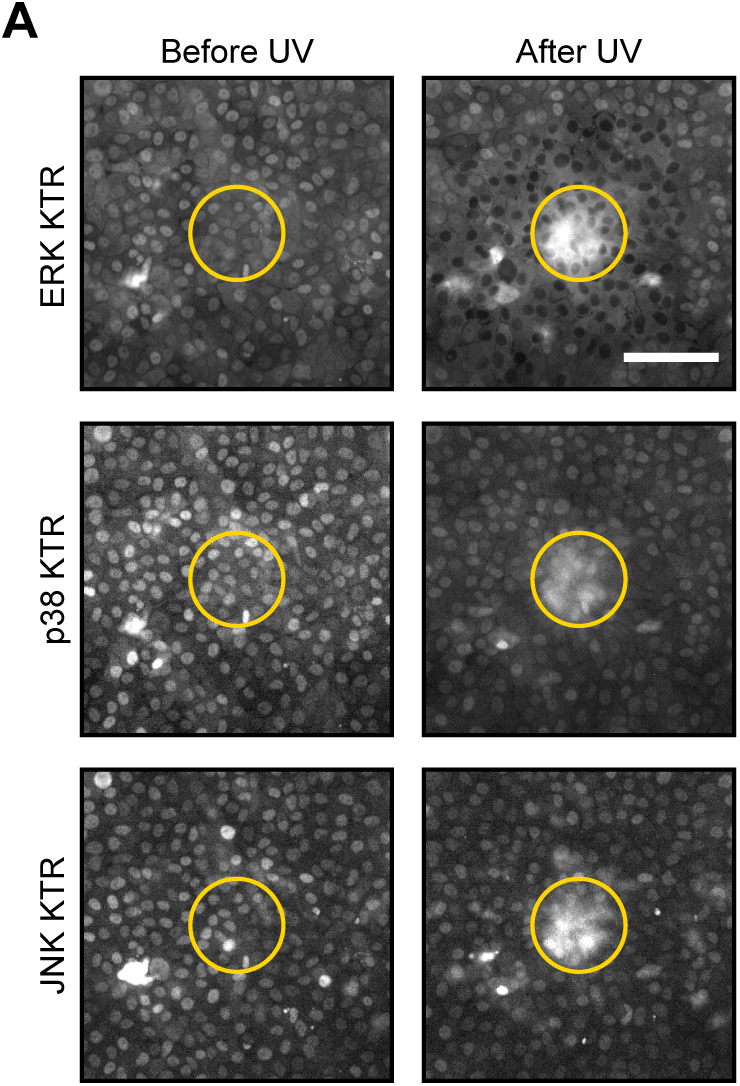
UV illumination leads to ERK activation of neighboring cells. **A**. Representative images of a cell line containing ERK KTR-mCer3, p38 KTR-mClov, and JNK KTR-mRuby2 before and after regional UV exposure (yellow circle) as in 5A. Scale bar = 100µm.

**Supplementary Figure 5.**
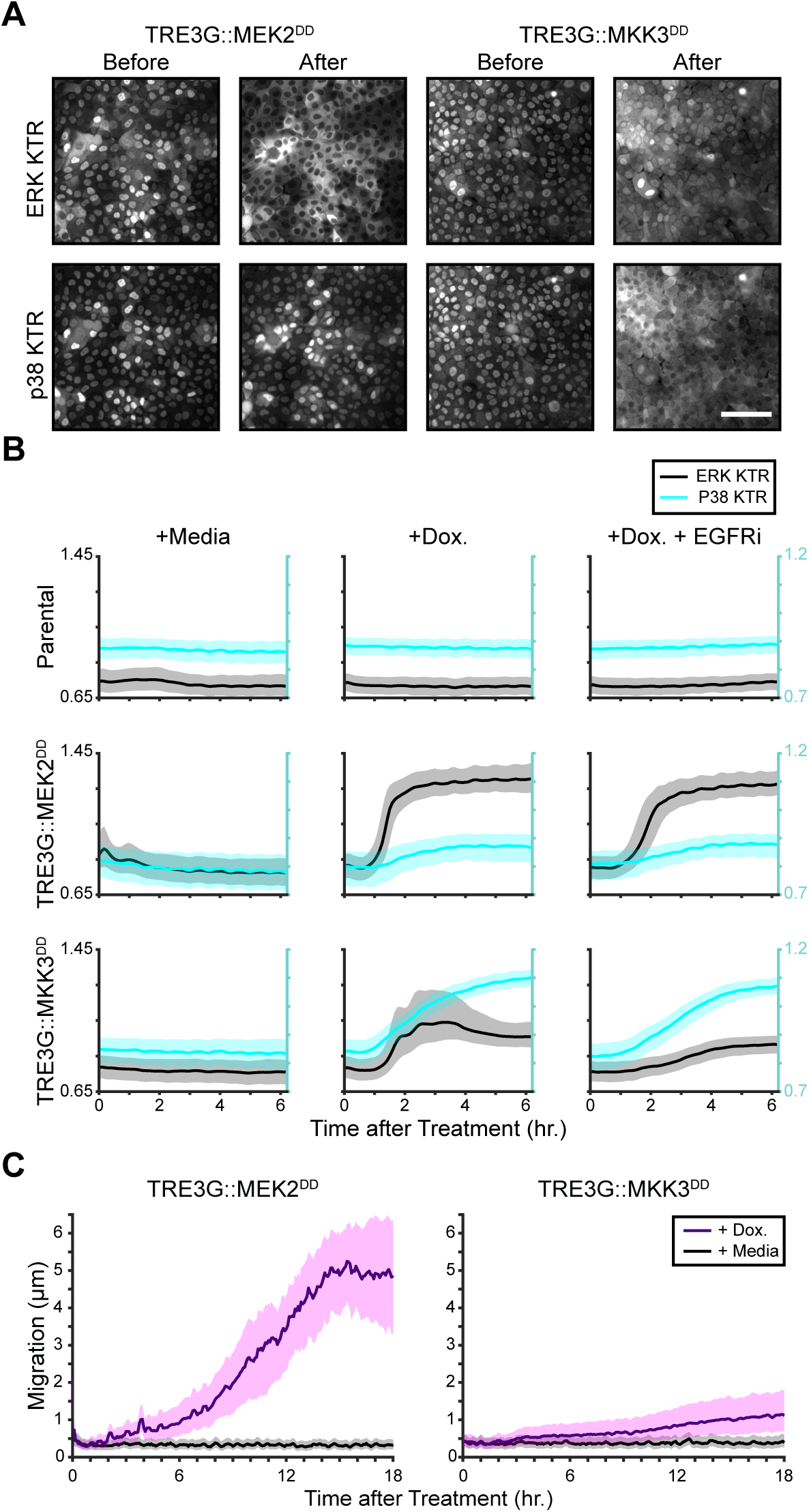
MAPK specificity of MEK2^DD^ and MKK3^DD^. **A**. Representative images from inducible MEK2^DD^ and MKK3^DD^ cells expressing H2B-iRFP, ERK KTR-mCer3 and p38 KTR-mClov before and after induction with doxycycline (2µg/ml). Scale bar = 100µm. **B**. Population average traces of ERK (black) and p38 (cyan) activity for indicated parental and inducible ERK/p38 KTR cells from A. Cocultures were treated with media, doxycycline, or doxycycline with EGFR inhibitor (5µM gefitinib) to limit paracrine ERK activation. Population averages and 25th-75th percentiles (shaded regions) are shown for n > 2000 cells. **C**. Migration of inducible cells from B plotted as µm change for every 5 min timepoint.

**Supplementary Figure 6.**
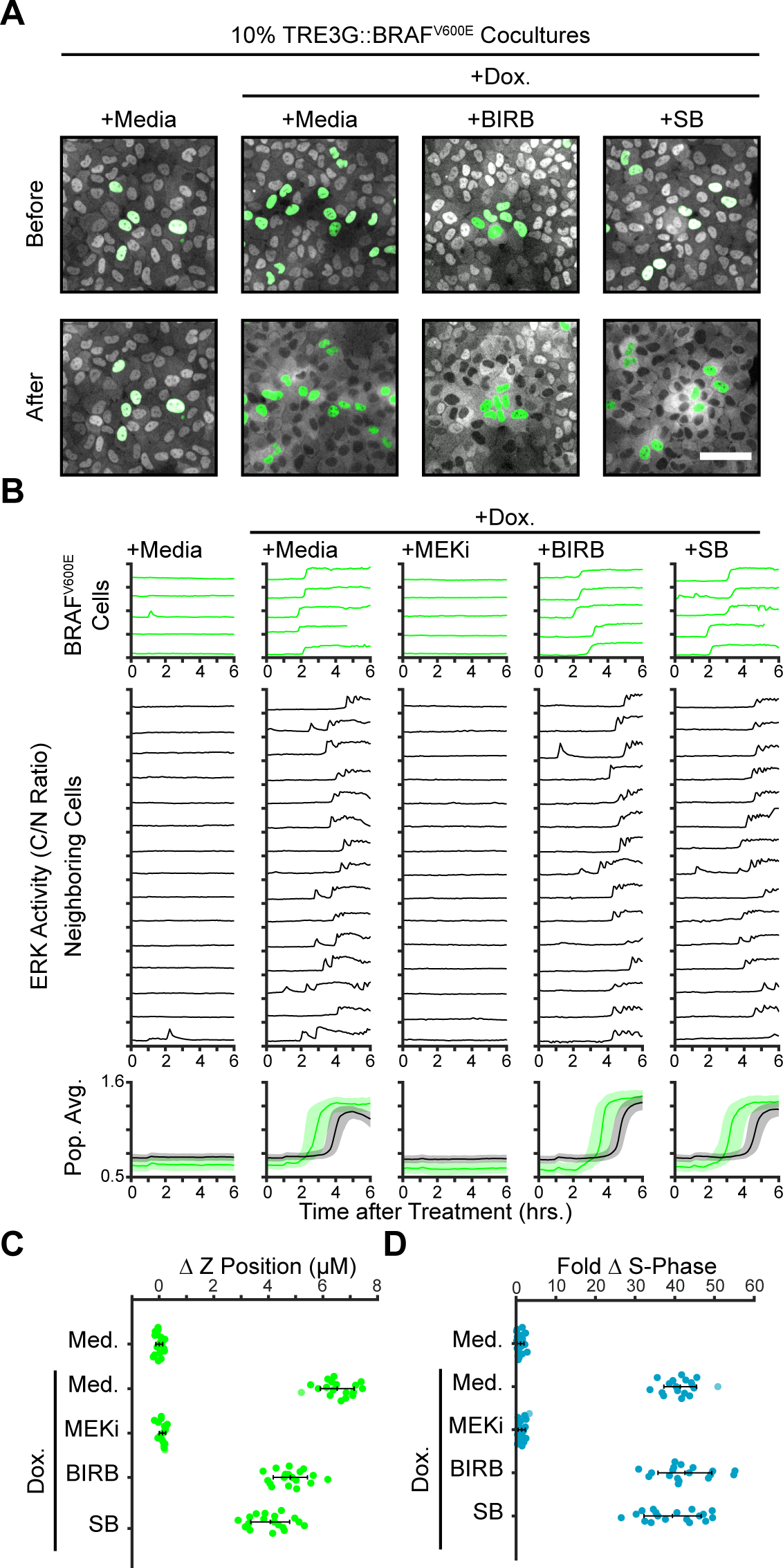
Oncogene-induced ERK signaling and cell behaviors are not p38-dependent. **A**. Representative images from 10% TRE3G::BRAF^V600E^ cocultures plated as in 2B before and after treatment with media or doxycycline (2µg/ml) alone or with p38 inhibitors (5µM BIRB 796 or 25µM SB 203580). Scale bar = 100µm. **B**. ERK activity traces, averages and 25th-75th percentiles from inducible BRAF^V600E^ cocultures from a represented as in 2C for n > 1600 cells. **C**. Extrusion of BRAF^V600E^ inducible cells from 10% cocultures treated as indicated and presented as in 3E for n=18 observations. **D**. EdU incorporation in 10% BRAF^V600E^ cocultures treated as indicated and presented as in Fig. 3H for n=18 observations (see methods).

**Supplementary Figure 7.**
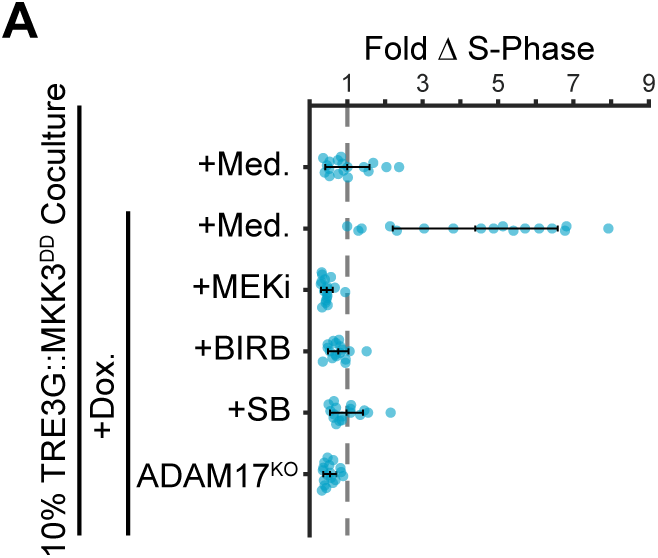
Mosaic p38 activation leads to ADAM17-EGFR dependent proliferation of neighboring cells. **A**. EdU of 10% MKK3^DD^ inducible cell cocultures treated for 24 hours with vehicle (+Media) or doxycycline (2µg/ml) in the presence of MEK inhibitor (MEKi, 5µM PD 0325901), p38 inhibitiors (5µM BIRB 796 or 25µM SB 203580) or in ADAM17^KO^ cells. Data presented as in 4B for n > 16 observations. See methods for details about EdU incorporation.

**Supplementary Figure 8.**
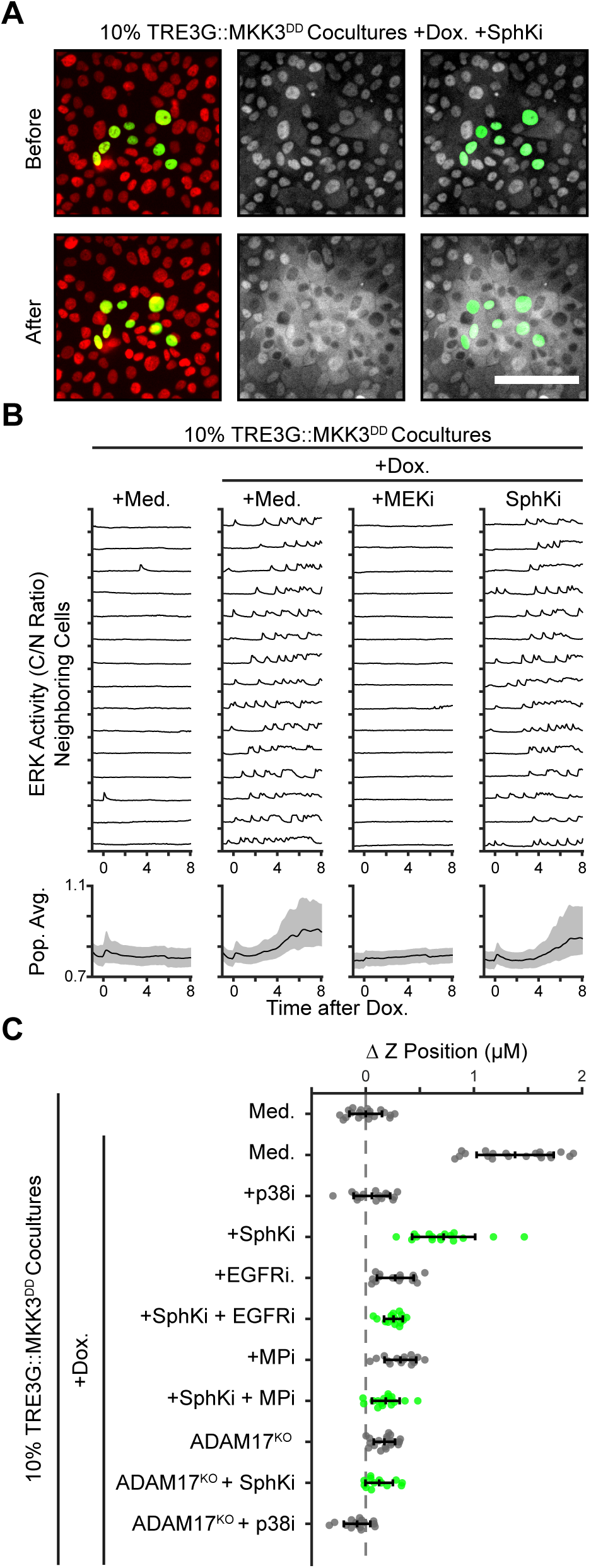
Shingosine-1-Phosphate is not required for signaling or extrusion effects of ADAM17-EGFR-ERK pathway. **A**. Representative images from 10% MKK3^DD^ cocultures treated with doxycycline (2µg/ml) and Sphingosine Kinase inhibitor (5µM SKII) presented as in 2B. Scale bar = 100µm. **B**. ERK activity traces of neighboring cells from MKK3^DD^ cocultures treated with media or doxycycline and Sphingosine Kinase or MEK inhibition presented as in 5H. Average traces and 25th-75th percentiles (shaded region) shown for neighboring cells on bottom for populations of n > 2000 cells. **C**. Extrusion analysis of 10% MKK3^DD^ cocultures presented as in 5I. Conditions in grey are reproduced from 5I for comparison with Sphingosine Kinase inhibitor and additive conditions (green).

**Supplementary Table 1.**

| Secreted Factor | +Dox. / -Dox. |  | WT / ADAM17-KO |  |
| --- | --- | --- | --- | --- |
|  | Fold Change | P-value | Fold Change | P-value |
| AREG | 8.35 | 5.88E-03 | 10.75 | 4.39E-03 |
| SDC4 | 5.32 | 9.18E-03 | 4.52 | 3.90E-03 |
| NECTIN1 | 3.67 | 1.49E-03 | 4.67 | 3.92E-03 |
| APP | 3.25 | 4.29E-03 | 1.98 | 1.03E-02 |
| PTPRS | 2.83 | 1.42E-02 | 3.23 | 2.18E-03 |
| PIK3IP1 | 2.83 | 2.89E-02 | 1.80 | 4.12E-02 |
| JAG1 | 2.75 | 1.24E-02 | 2.85 | 2.28E-03 |
| DDR1 | 2.69 | 2.37E-02 | 2.36 | 7.74E-03 |
| DAG1 | 2.67 | 1.73E-02 | 2.16 | 1.27E-02 |
| SFRP1 | 2.60 | 3.75E-02 | 2.70 | 3.69E-02 |
| NEO1 | 2.53 | 1.02E-02 | 2.56 | 8.18E-03 |
| MDK | 2.51 | 3.16E-02 | 1.98 | 1.54E-02 |
| HLA-C | 2.48 | 1.62E-02 | 1.99 | 2.17E-02 |
| MET | 2.22 | 3.56E-02 | 2.21 | 1.60E-02 |
| PTPRF | 2.20 | 1.10E-02 | 2.36 | 1.87E-03 |
| SDC1 | 2.16 | 3.73E-02 | 1.61 | 4.11E-02 |
| MSLN | 2.15 | 1.28E-02 | 1.59 | 4.60E-02 |
| SPINT1 | 2.10 | 1.93E-02 | 1.83 | 1.90E-02 |
| HLA-B | 2.07 | 1.23E-02 | 2.87 | 9.49E-03 |
| PTK7 | 2.05 | 3.81E-02 | 3.37 | 3.25E-03 |
| APLP2 | 1.91 | 7.05E-03 | 1.93 | 1.47E-03 |
| PROCR | 1.87 | 4.70E-02 | 2.96 | 4.26E-02 |
| HLA-A | 1.71 | 2.05E-02 | 2.22 | 1.05E-02 |
| CCT3 | 1.54 | 1.51E-03 | 1.65 | 1.07E-02 |

